# Abundance of Enterovirus C in RD-L20B cell culture negative stool samples from Acute Flaccid Paralysis cases in Nigeria is geographically defined

**DOI:** 10.1101/222935

**Authors:** E Donbraye, O.I. Olasunkanmi, B.A. Opabode, T.R. Ishola, T.O.C. Faleye, M.O. Adewumi, J.A. Adeniji

**Author notes:** To whom all correspondence should be addressed: AUTHOR NAME and EMAIL. DONBRAYE, Emmanuel;, OLASUNKANMI, Oluwatayo Israel, OPABODE, Babatunde Ayoola, ISHOLA, Temitayo Racheal, FALEYE, Temitope Oluwasegun Cephas, ADEWUMI, Moses Olubusuyi, ADENIJI, Johnson Adekunle.

## Abstract

We recently showed that Enteroviruses (EVs); majorly species Cs (EV-Cs) were present in about 46.7% of faecal samples from children <15 years old diagnosed with Acute Flaccid Paralysis (AFP) in Nigeria but declared to be EV free by the RD-L20B cell culture based algorithm. In this study, we investigated whether this observed preponderance of EVs (and EV-Cs) in such samples vary by geographical region.

In all, 108 samples (i.e. 54 paired stool suspensions from 54 AFP cases) previously confirmed negative for EVs by the WHO recommended RD-L20B cell culture based algorithm were analyzed in this study. The 108 samples were made into 54 pools (27 each from Northwest [NW] and Southsouth [SS] Nigeria). All samples were subjected to RNA extraction, cDNA synthesis and the WHO recommended seminestedPCR (snPCR) assay and its modifications. All amplicons were sequenced, and enteroviruses identified using the enterovirus genotyping tool and phylogenetic analysis.

Altogether, EVs were detected in 16 (29.63%) of the 54 samples screened but successfully identified in 14 (25.93%): 10 from NW- and 4 from SS-Nigeria. Precisely, one (7.14%), two (14.29%) and 11 (78.57%) of the strains detected were EV-A, EV-B and EV-C respectively. The 10 strains from NW-Nigeria are 7 EV types and include CV-A10, E29, CV-A13, CV-A17, CV-A19, CV-A24 and EV-C99. The four EV types recovered from SS-Nigeria include E31, CV-A1, EV-C99 and EV-C116. EV-C99 is the only EV type that was detected in both NW- and SS-Nigeria.

The results of this study showed that the preponderance of EVs and consequently EV-Cs in AFP samples declared to be EV free by the RD-L20B cell culture based algorithm vary by geographical region in Nigeria. It further confirmed the EV-B bias of the RD-L20B cell culture based algorithm.

## Introduction

Enteroviruses are members of the genus *Enterovirus* in the family *Picornaviridae*, order *Picornavirales*. There are 13 species in the genus and the best studied member of the genus, poliovirus, belongs to species C (EV-C) (www.picornaviridae.com). Courtesy the resolution of the World Health Assembly (WHA) to eradicate poliovirus (WHO, 1988) the Global Polio Eradication Initiative (GPEI) has eliminated indigenous strains of the virus globally except in three countries; Afghanistan, Nigeria and Pakistan (www.polioeradication.org). This eradication effort is driven by a two-pronged system; vaccination and surveillance. Vaccination uses both an Inactivated Polio Vaccine (IPV) and the attenuated live virus vaccine (Oral Polio Vaccine [OPV]) to inhibit virus transmission while surveillance looks for both the street (wild) virus and attenuated live (vaccine) virus in both children with acute flaccid paralysis (AFP) and the environment (WHO, 2015).

Ordinarily, subsequent to large scale co-ordinated immunization activities, the attenuated vaccine virus should stop circulating in the population in about eight weeks (Nathanson and Kew, 2010). It has however, been observed that many times this does not happen because the virus easily regains its transmissibility and pathogenicity by reverting to wild type genotype and consequently, phenotype (Combelas et al., 2011). Such wild type poliovirus strains of vaccine origin are referred to as Vaccine Derived Polioviruses (VDPV) (Burns et al., 2014). Many VDPVs (as exemplified in the strains that caused the outbreak that started in Nigeria in 2005 and lasted about a decade [Wassilak et al., 2011, Burns et al., 2013; 2014]), are recombinant in nature. These virus strains have a structural region of OPV origin and a non-structural region of non-polio enterovirus species C (NPEV-C) origin (OPV/NPEV-C).

This role of NPEV-Cs in VDPV emergence exemplifies their importance, and the need to carefully catalogue their diversity and geographical distribution. It has however been shown (Sadeuh-Mba 2013, Adeniji and Faleye, 2014b, Faleye and Adeniji, 2015b, Adeniji et al., 2017) that the RD-L20B cell culture based algorithm (WHO, 2003; 2004) recommended for and largely in use by the Global Polio Laboratory Network (GPLN) under-reports the preponderance of EV-Cs. This is because the L20B cell line component of the algorithm is largely specific for poliovirus detection due to its expression of the human poliovirus receptor (Pipkin et al. 1993),while the RD cell line component that is supposed to be more promiscuous for EV detection is EV-B bias (Sadeuh-Mba 2013, Adeniji and Faleye, 2014a, Faleye and Adeniji, 2015a & b, Adeniji et al., 2017)

We recently showed (Adeniji et al., 2017) that EVs were present in about 46.7% of faecal samples from children <15 years old diagnosed with AFP but declared to be EV free by the RD-L20B cell culture based algorithm (WHO, 2003, 2004). In addition, we showed that majority of the EVs recovered in these samples were EV-Cs. This suggests that the EV-B bias of the RD component of the RD-L20B cell culture based algorithm (WHO, 2003, 2004) selectively accumulated as ‘negatives’, samples that lacked EV-Bs but possibly had EV-Cs.

Nigeria has a history of varying regional peculiarities that significantly influence the preponderance and dynamics of polioviruses (CDC, 2006& 2010; Mach et al., 2014; DuintjerTebbens et al., 2015, Faleye and Adeniji, 2015a). For example, of the 23 lineages of circulating VDPV2s (cVDPV2) that emerged in Nigeria between 2005 and 2011 (Burns et al., 2013), 12 (52%) and none (0%) emerged in Northwest (NW) and Southsouth (SS) Nigeria, respectively. Hence, in this study, we investigated whether the observed preponderance of EVs (and EV-Cs) inAFP samples declared to be EV free by the RD-L20B cell culture based algorithmin the country vary by region.

## Methodology

### Sample Collection and Processing

In August 2015, 86.26% (747/866) of the AFP cases received by the WHO accredited polio laboratory in Ibadan, Nigeria (subsequently referred to as the polio lab) were declared negative for enteroviruses using the WHO recommended RD-L20B cell culture based algorithm (WHO, 2004). In this study, we randomly selected 54 (27 each from North-West and South-South Nigeria [Figure 1]) of these cases (i.e 7.23% [54/747]) for further analysis. All the pairs of stool suspensions made from these cases were collected from the archives of the polio lab and pooled. That is, the 108 stool suspensions from the 54 cases were pooled into 54 stool suspensions and subsequently analyzed in this study.

**Figure 1:**
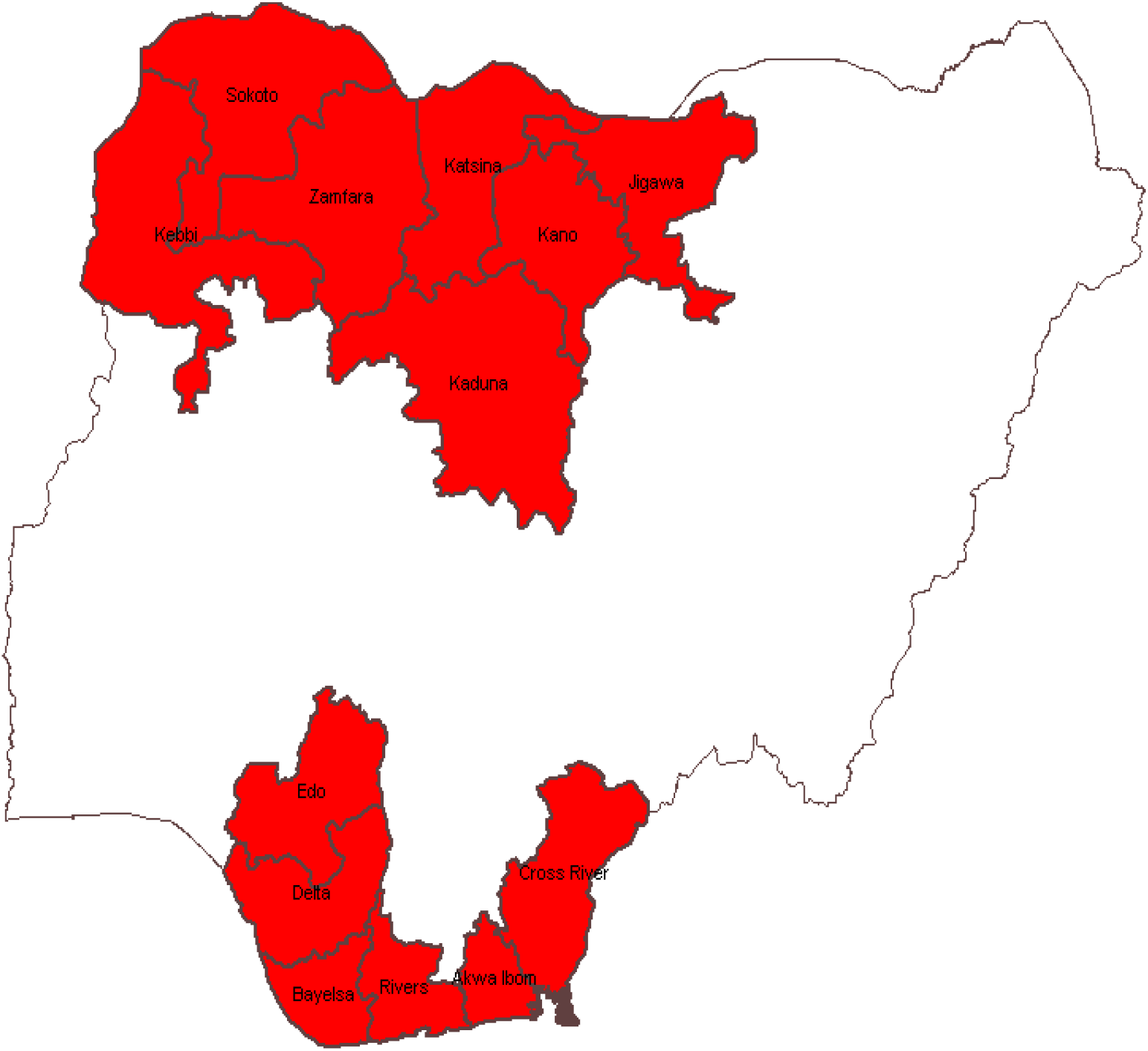
The Map of Nigeria highlighting North-West (7 states in the top left hand corner) and South-South (6 states in the bottom of the image) Nigeria in Red.

### RNA Extraction, cDNA Synthesis and seminested Polymerase Chain Reaction (snPCR)

RNA extraction, cDNA synthesis and snPCR were done as recently described (Faleye et al., 2016; Adeniji et al., 2017). Briefly, RNA extraction and cDNA synthesis were done using Jena Bioscience kits for RNA extraction and cDNA synthesis kits (Jena Bioscience, Jena, Germany), respectively, according to manufacturer’s recommendations. However, in place of random hexamers, primers AN32-AN35 (Nix et al., 2006, WHO, 2015) were used for cDNA synthesis.

For the snPCR assay, there was one first round and three (Panenterovirus [PE], Enterovirus Species A or C [EV-A/C] and Enterovirus Species B [EV-B]) second round PCR assays. All assays were done in 30μL volumes. Each contained 6μL of Red Load Taq, and 0.3μL each of the forward and reverse primers. For the first round PCR assay, 13.4μL of RNase free water and 10μL of cDNA were also added to the mix. While, for the second round PCR assay, 20.4μL of RNase free water and 3μL of first round PCR product were added to the mix. The primers for the first round PCR were 224 and 222 (Nix et al, 2006). For the second round PCR assay, all three assays used reverse primer AN88 while the forward primers were AN89, 189 and 187 for the PE, EV-A/C and EV-B PCR assays, respectively. The cycling conditions for the assays were similar but for the extension time that was 60 and 30 seconds respectively for the first and second round PCR assays, respectively. All PCR products were resolved on 2% agarose gels stained with ethidium bromide and viewed using a UV transilluminator. Samples in which the second round PCR assay successfully amplified the expected ~350bp amplicon, were considered positive

### Amplicon Sequencing and Enterovirus Identification

All the appropriately sized amplicons (~350bp) generated from the second round PCR assays were shipped to MacrogenInc, Seoul, South Korea, for purification and nucleotide sequencing. Subsequently, the enterovirus species and genotypes were determined using the enterovirus genotyping tool (EGT) (Kroneman et al., 2011).

### Phylogenetic Analysis

Being the most detected, only the EV-Cs were subjected to phylogenetic analysis. The CLUSTAL W program in MEGA 5 software (Tamura et al., 2011) was used for multiple sequence alignments with default settings. Afterwards, neighbor-joining trees were constructed using the Kimura-2 parameter model (Kimura, 1980) and 1,000 bootstrap replicates in the same MEGA 5 software.

### Nucleotide Sequence Accession Numbers

The sequences generated in this study have been deposited in GenBank under accession numbers MG252505-MG252518.

## RESULTS

### North-West Nigeria PCR Result

Eleven (40.74%) of the 27 samples from NW Nigeria were positive (i.e. with the expected ~350bp successfully amplified) for the PE PCR assay. Ten (37.04%) of the 27 samples were positive (i.e. with the expected ~350bp successfully amplified) for the EV-A/C PCR assay. Nine of the 10 samples positive for the EV-A/C PCR assay, were also positive for the PE PCR assay. Specifically, one (sample 13) of the 11 samples positive for the PE PCR assay was negative for the EV-A/C PCR assay and one (sample 23) of the 10 samples positive for the EV-A/C PCR assay was negative for the PE PCR assay.

Only two (7.4%) of the 27 samples were positive for the EV-B PCR assay. These two samples were also positive for the PE PCR assay and one of them (sample 2) was positive for the EV-A/C PCR assay too. Thus, 9 (81.82%) of the 11 samples positive for the PE PCR assay were negative for the EV-B PCR assay. Ultimately, 12 (44.44%) of the 27 samples were positive for at least one of the three assays but only one sample (Sample 2) was positive for all three assays (Table 1).

**Table 1:** Results of RT-snPCR screen of cell culture negative AFP stool samples from NW-Nigeria.

| S/N | Lab ID | RT-snPCR RESULT & ENTEROVIRUS ID |  |  | SUMMARY | SPECIES |
| --- | --- | --- | --- | --- | --- | --- |
|  |  | PE | EV-A/C | EV-B |  |  |
| 1 | 2 | + | + | + | EV-C99 | EV-C |
| 2 | 7 | + | + | --- | CV-A19 | EV-C |
| 3 | 9 | + | + | --- | CV-A24 | EV-C |
| 4 | 11 | + | --- | + | E-29 | EV-B |
| 5 | 13* | + | --- | --- |  |  |
| 6 | 14 | + | + | --- | EV-C99 | EV-C |
| 7 | 15 | + | + | --- | CV-A17 | EV-C |
| 8 | 16 | + | + | --- | CV-A17 | EV-C |
| 9 | 19 | + | + | --- | EV-C99 | EV-C |
| 10 | 21 | + | + | --- | CV-A10 | EV-A |
| 11 | 23* | --- | + | --- |  |  |
| 12 | 24 | + | + | --- | CV-A13 | EV-C |
| TOTAL |  | 11 | 10 | 2 | 10 |  |

\* = Sequence data not usable due to multiple peaks
= Positive for the assay
= Negative for the assay

### South-South Nigeria PCR Result

Four (14.82%) of the 27 samples from SS Nigeria were positive for the PE PCR assay. Two (7.4%) of the 27 samples were positive for the EV-A/C PCR assay. These two samples were also positive for the PE PCR assay. Thus, two (50.0%) of the four samples positive for the PE PCR assay were negative for the EV-A/C PCR assay. Only one (3.7%) of the 27 samples was positive for the EV-B PCR assay. This sample was also positive for the PE PCR assay. Hence, three (75.0%) of the four samples positive for the PE PCR assay were negative for the EV-B PCR assay. Ultimately, 4 (14.82%) of the 27 samples from SS Nigeria were positive for at least one of the three assays but none of the samples were positive for all three assays (Table 2).

**Table 2:** Results of RT-snPCR screen of cell culture negative AFP stool samples from SS-Nigeria.

| S/N | Lab ID | RT-snPCR RESULT &<br>ENTEROVIRUS ID |  |  | SUMMARY | SPECIES |
| --- | --- | --- | --- | --- | --- | --- |
|  |  | PAN-ENTERO | EV-A/C | EV-B |  |  |
| 1 | 9 | + | + | --- | EV-C116 | EV-C |
| 2 | 16 | + | --- | + | E-31 | EV-B |
| 3 | 20 | + | --- | --- | EV-C99 | EV-C |
| 4 | 26 | + | + | --- | CV-A1 | EV-C |
| TOTAL |  | 4 | 2 | 1 | 4 |  |

+ = Positive for the assay
--- = Negative for the assay

### North-West Nigeria Sequencing and Genotyping Results

All the 11 amplicons generated by the PE PCR assay were successfully sequenced. However, the sequence data for one of the amplicons (Sample 13) was not usable due to multiple peaks. Strains identified were Enterovirus (EV) C99 (3 isolates), Echovirus-29 (E29) (1 isolate), Coxsackievirus (CV) A10 (1 isolate), CV-A13 (1 isolate), CV-A17 (2 isolates), CV-A19 (1 isolate) and CV-A24 (1 isolate). Of the 10 strains successfully identified via the PE PCR assay, one (CV-A10), one (E29) and eight belong to EV-A, EV-B and EV-C, respectively (Table 1).

For the EV-A/C assay, all the 10 amplicons generated were successfully sequenced. However, the sequence data for one of the amplicons (Sample 23) was not usable due to multiple peaks. Strains identified were EV-C99 (3 isolates), CV-A10 (1 isolate), CV-A13 (1 isolate), CV-A17 (2 isolates), CV-A19 (1 isolate) and CV-A24 (1 isolate). Of the nine (9) strains successfully identified via the EV-A/C assay, one (CV-A10) and eight belong to EV-A and EV-C, respectively (Table 1).

Only one (sample 2) of the two amplicons generated by the EV-B assay was sequenced and identified as EV-C99.The second amplicon (Sample 11) was not sequenced because the intensity of the band was too weak (Table 1). In all, EVs were detected in 12 (44.44%) of the 27 samples screened but successfully identified in 10 (37.04%) of the samples.

### South-South Nigeria Sequencing and Genotyping Results

All the 4 amplicons generated by the PE PCR assay were successfully sequenced and three of the strains were identified as CV-A1 (1 isolate), E31 (1 isolate) and EV-C99 (1 isolate). The fourth strain was identified as a member of EV-C but the type was unassigned by the EGT. However, the EGT phylogenetic tree showed it to cluster with EV-C116 but with a bootstrap support of 67 (supplementary Figure 1). Hence, it will be here on referred to as EV-C116. Of the four (4) strains successfully identified via the PE PCR assay, one (E31) and three belong to EV-B and EV-C, respectively (Table 2). Both amplicons generated by the EV-A/C PCR assay were successfully sequenced and the strains identified as CV-A1 (1 isolate) and EV-C116 (1 isolate). The EV-C116 was also typed using the EGT phylogenetic tree. Both strains successfully identified via the EV-A/C assay belong to EV-C (Table 2).The only amplicons generated by the EV-B PCR assay was sequenced and identified as E31. This strain belongs to EV-B (Table 2). In all, EVs were detected and successfully identified in four (14.81%) of the 27 samples screened.

### North-West and South-South Nigeria Sequencing and Genotyping Result summary

Altogether, EVs were detected in 16 (29.63%) of the 54 samples screened but successfully identified in 14 (25.93%) samples. Precisely, one (7.14%), two (14.29%) and 11 (78.57%) of the strains detected were EV-A, EV-B and EV-C respectively.

### Comparison of EV types detected in the two Geo-Political Regions

The seven EV types recovered from NW-Nigeria spread over three EV Species (EV-A, B and C) and include CV-A10 (EV-A), E29 (EV-B) and CV-A13, CV-A17, CV-A19, CV-A24 and EV-C99 (EV-Cs). On the other hand, the four EV types recovered from SS-Nigeria spread over two EV Species (EV-B and C). These are E31 (EV-B) and CV-A1, EV-C99 and EV-C116 (EV-Cs). EV-C99 (an EV-C) is the only EV type that was detected in both NW- and SS-Nigeria (Table 3).

**Table 3:** Diversity of Enteroviruses recovered from the two geopolitical regions analyzed in this study.

| S/N | NW-<br>NIGERIA | NUMBER OF<br>ENTEROVIRUS<br>TYPES | SS-<br>NIGERIA | NUMBER OF<br>ENTEROVIRUS<br>TYPES | TOTAL NUMBER<br>OF ENTEROVIRUS<br>TYPES |
| --- | --- | --- | --- | --- | --- |
| EV-A | CV-A10 | 1 | Nil | 0 | 1 |
| EV-B | E-29 | 1 | E-31 | 1 | 2 |
| EV-C | CV-A13,<br>CV-A17,<br>CV-A19,<br>CV-A24,<br>EV-C99* | 5 | CV-A1,<br>EV-C99*,<br>EV-C116 | 3 | 7 |
| TOTAL |  | 7 |  | 4 | 10* |

\* = Though EV-C99 was recovered from both regions, it is calculated as the same EV type. Hence the reason the total number of EV types is 10 and not 11.

### Phylogenetic Analysis

Being the most detected, only the seven (7) EV-C types found in this study were further subjected to phylogenetic analysis. The CV-A1 strains found in GenBank alongside the one described in this study grouped into five genotypes based on bootstrap values (Figure 2). The CV-A1 detected in this study grouped with genotype B and is most similar to the CV-A1 recently recovered from a child with AFP in Nigeria in 2015. The CV-A1 strain detected in this study is different from the strain recovered from a healthy child in 2014 in Nigeria. The 2014 strain belongs to genotype E (Figure 2). The Nigerian strains of CV-A19 and EV-C116 are being described for the first time in this study and based on the topology of the tree (Figure 2), they seem to be distantly related to the strains available in GenBank.

**Figure 2:**
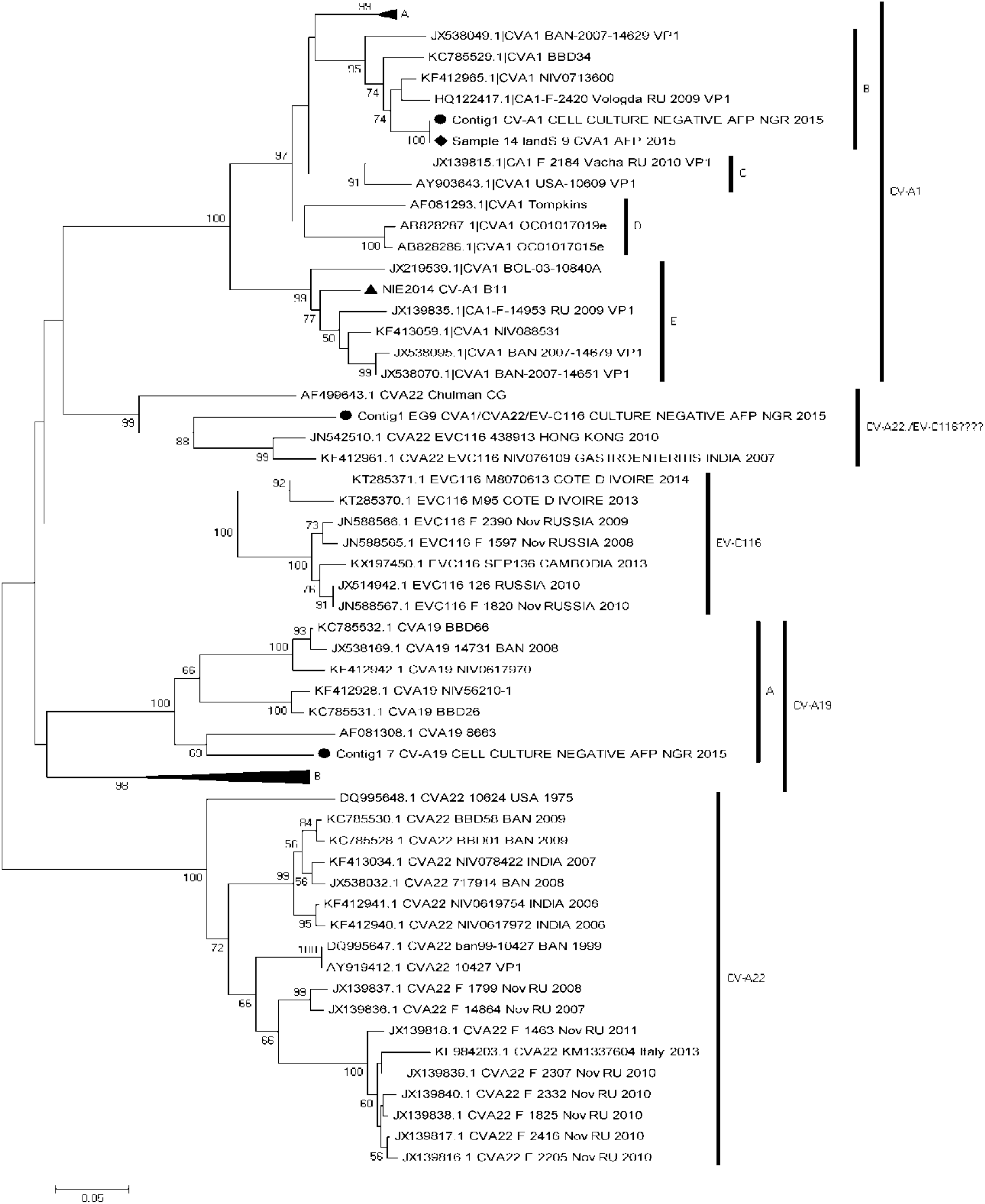
Phylogram of genetic relationship between VP1 nucleotide sequences of CV-A1, CV-A19, CV-A22 and EV-C116 strains. The phylogenetic tree is based on an alignment of the partial VP1 sequences. The newly sequenced strains are indicated with black circle. Strains previously detected in Nigeria are indicated with black diamond (2015) and triangle (2014).Bootstrap values are indicated if >50%.

The EV strain identified as EV-C116 using the EGT phylogenetic tree (supplementary figure 1) did not cluster with EV-C116 strains recovered from GenBank. Rather it clustered with reference CV-A22 strain Chulman and two other CV-A22 strains (Figure 2). To resolve this inconsistency, a BLASTn search of the sequence data of this EV strain was done. The BLASTn result suggested that it was only similar to CV-A22 strain recovered in Hong Kong in 2010 and this was consistent with the results of our similarity analysis (Table 4).

**Table 4:** Similarity analysis of Contig1 EG9 CVA1/CVA22/EV-C116 CULTURE NEGATIVE AFP NGR 2015 with reference strains downloaded from GenBank. Only strains with similarity more than 60% are shown.

| Species 1 | Species 2 | Dist % | Siml% |
| --- | --- | --- | --- |
| Contig1 EG9 CVA1/CVA22/EV-C116 CULTURE NEGATIVE AFP NGR 2015 | KF412961.1 CVA22 EVC116 NIV076109 GASTROENTERITIS INDIA 2007 | 16.02858 | 83.97142 |
| Contig1 EG9 CVA1/CVA22/EV-C116 CULTURE NEGATIVE AFP NGR 2015 | JN542510.1 CVA22 EVC116 438913 HONG KONG 2010 | 19.38896 | 80.61104 |
| Contig1 EG9 CVA1/CVA22/EV-C116 CULTURE NEGATIVE AFP NGR 2015 | AF499643.1 CVA22 Chulman CG | 21.26653 | 78.73347 |
| Contig1 EG9 CVA1/CVA22/EV-C116 CULTURE NEGATIVE AFP NGR 2015 | KF412928.1 CVA19 NIV56210-1 | 35.88158 | 64.11842 |
| Contig1 EG9 CVA1/CVA22/EV-C116 CULTURE NEGATIVE AFP NGR 2015 | KF413059.1\|CVA1 NIV088531 | 36.52963 | 63.47037 |
| Contig1 EG9 CVA1/CVA22/EV-C116 CULTURE NEGATIVE AFP NGR 2015 | JX174176.1\|CVA1 HT-THLH02F-XJ-CHN-2011 | 37.11544 | 62.88456 |
| Contig1 EG9 CVA1/CVA22/EV-C116 CULTURE NEGATIVE AFP NGR 2015 | JX181912.1\|CVA1-HG2010038 | 37.13184 | 62.86816 |
| Contig1 EG9 CVA1/CVA22/EV-C116 CULTURE NEGATIVE AFP NGR 2015 | JX181911.1\|CVA1-HG2010043 | 37.13184 | 62.86816 |
| Contig1 EG9 CVA1/CVA22/EV-C116 CULTURE NEGATIVE AFP NGR 2015 | JX181910.1\|CVA1-HG2011009 | 37.13184 | 62.86816 |
| Contig1 EG9 CVA1/CVA22/EV-C116 CULTURE NEGATIVE AFP NGR 2015 | KC785531.1 CVA19 BBD26 | 37.96521 | 62.03479 |
| Contig1 EG9 CVA1/CVA22/EV-C116 CULTURE NEGATIVE AFP NGR 2015 | JX174177.1\|CVA1 KS-ZPH01F-XJ-CHN-2011 | 38.46617 | 61.53383 |
| Contig1 EG9 CVA1/CVA22/EV-C116 CULTURE NEGATIVE AFP NGR 2015 | KT285370.1 EVC116 M95 COTE D IVOIRE 2013 | 38.62784 | 61.37216 |
| Contig1 EG9 CVA1/CVA22/EV-C116 CULTURE NEGATIVE AFP NGR 2015 | AB828287.1\|CVA1 OC01017019e | 39.25287 | 60.74713 |
| Contig1 EG9 CVA1/CVA22/EV-C116 CULTURE NEGATIVE AFP NGR 2015 | AB828286.1\|CVA1 OC01017015e | 39.25287 | 60.74713 |
| Contig1 EG9 CVA1/CVA22/EV-C116 CULTURE NEGATIVE AFP NGR 2015 | KY865976.1 CVA19 3101300403 2 2013 CEVS | 39.29682 | 60.70318 |
| Contig1 EG9 CVA1/CVA22/EV-C116 CULTURE NEGATIVE AFP NGR 2015 | KY865975.1 CVA19 3101300402 2013 CEVS | 39.29682 | 60.70318 |
| Contig1 EG9 CVA1/CVA22/EV-C116 CULTURE NEGATIVE AFP NGR 2015 | KY865822.1 CVA19 3101300163 2013 CEVS | 39.29682 | 60.70318 |
| Contig1 EG9 CVA1/CVA22/EV-C116 CULTURE NEGATIVE AFP NGR 2015 | KY865821.1 CVA19 3101300162 2013 CEVS | 39.29682 | 60.70318 |
| Contig1 EG9 CVA1/CVA22/EV-C116 CULTURE NEGATIVE AFP NGR 2015 | JX538070.1\|CVA1 BAN-2007-14651 VP1 | 39.34727 | 60.65273 |
| Contig1 EG9 CVA1/CVA22/EV-C116 CULTURE NEGATIVE AFP NGR 2015 | KT285371.1 EVC116 M8070613 COTE D IVOIRE 2014 | 39.40416 | 60.59584 |
| Contig1 EG9 CVA1/CVA22/EV-C116 CULTURE NEGATIVE AFP NGR 2015 | AF081293.1\|CVA1 Tompkins | 39.40416 | 60.59584 |
| Contig1 EG9 CVA1/CVA22/EV-C116 CULTURE NEGATIVE AFP NGR 2015 | JN588567.1 EVC116 F 1820 Nov RUSSIA 2010 | 39.46743 | 60.53257 |
| Contig1 EG9 CVA1/CVA22/EV-C116 CULTURE NEGATIVE AFP NGR 2015 | JX514942.1 EVC116 126 RUSSIA 2010 | 39.68199 | 60.31801 |
| Contig1 EG9 CVA1/CVA22/EV-C116 CULTURE NEGATIVE AFP NGR 2015 | KC785529.1\|CVA1 BBD34 | 39.89393 | 60.10607 |
| Contig1 EG9 CVA1/CVA22/EV-C116 CULTURE NEGATIVE AFP NGR 2015 | DQ455609.1\|CVA1 Germany-922-2005 | 39.89393 | 60.10607 |

The CV-A13 strain recovered in this study was from NW-Nigeria (Table 3). It was most similar to a CV-A13 strain also recovered from a child with AFP in Nigeria in 2015 (Figure 3). Both strains belong to the cluster sub-Saharan Africa 3 alongside strains previously described in Cameroon and Central Africa Republic (Figure 3).

**Figure 3:**
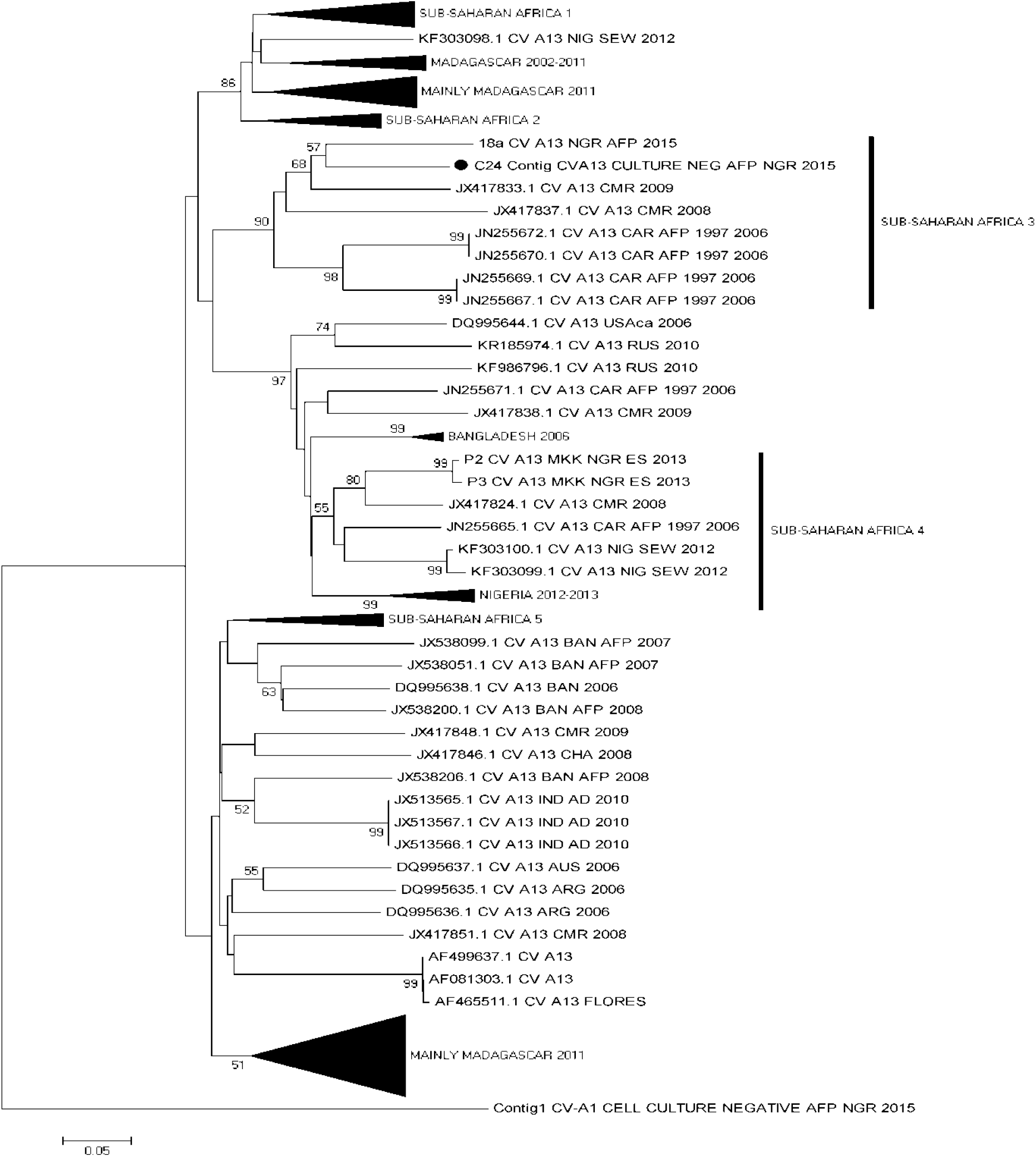
Phylogram of genetic relationship between VP1 nucleotide sequences of CV-A13 strains. The phylogenetic tree is based on an alignment of the partial VP1 sequences. The newly sequenced strain is indicated with a black circle. Bootstrap values are indicated if >50%. The labelled vertical bars are for ease of reference only.

The two CV-A17 strains described in this study were from NW-Nigeria (Table 3). Both strains belong to a cluster containing mainly strains from sub-Saharan Africa. They however share a common ancestor with a strain recovered in the Philippines in 2009 (Figure 4). The two CV-A17 strains described in this study are different from CV-A17 strains recovered in 2003 and 2015 from children with AFP in Nigeria (Figure 4).

**Figure 4:**
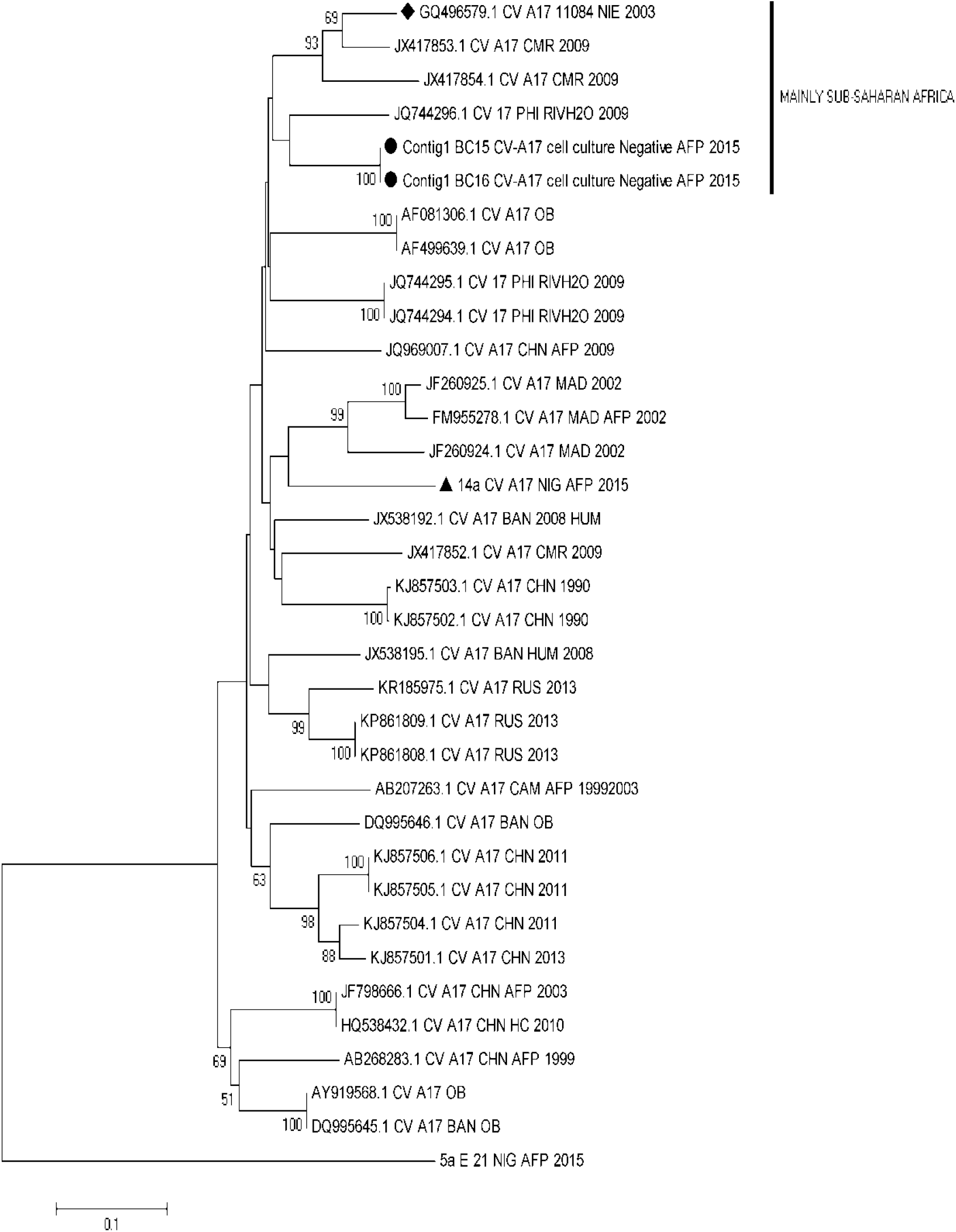
Phylogram of genetic relationship between VP1 nucleotide sequences of CV-A17 strains. The phylogenetic tree is based on an alignment of the partial VP1 sequences. The newly sequenced strain is indicated with a black circle. Strains recovered in Nigeria in 2003 and 2015 are indicated with a black diamond and triangle, respectively. Bootstrap values are indicated if >50%. The labelled vertical bars are for ease of reference only.

The CV-A24 strains described in this study belongs to a cluster of CV-A24 that has repeatedly been recovered from sub-Saharan Africa (Figure 5). It was however different from the strain recovered in 2010 from sewage contaminated water in Nigeria. Rather, the CV-A24 strain recovered in this study was most similar to a strain recovered in Cameroon (Figure 5).

**Figure 5:**
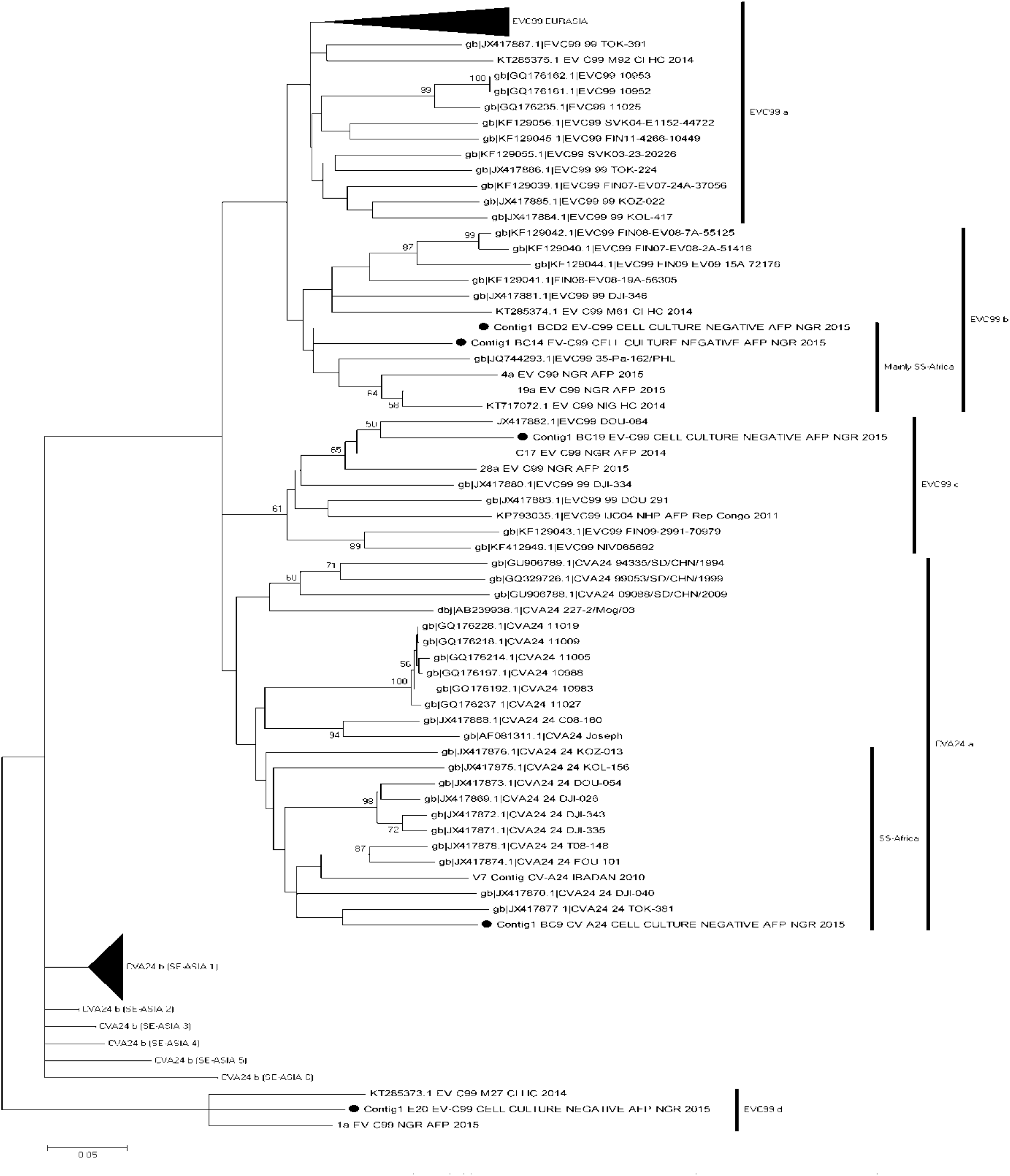
Phylogram of genetic relationship between VP1 nucleotide sequences of CV-A24 and EV-C99 strains. The phylogenetic tree is based on an alignment of the partial VP1 sequences. The newly sequenced strains are indicated with a black circle. Bootstrap values are indicated if >50%. The labelled vertical bars are for ease of reference only.

Four (one and three from SS- and NW-Nigeria, respectively) EV-C99 strains were recovered in this study. The strain recovered from SS-Nigeria (Figure 5; Cluster EV-C99 d) clustered with two strains. One of which was also recovered in 2015 from a child with AFP in Nigeria. The other strain was recovered in 2014 from a healthy child in Cote d’Ivore (Figure 5).

The three strains from NW-Nigeria fell into two clusters (EV-C99 b and c) (Figure 5). Two of the EV-C99 strains recovered in this study belong to cluster EV-C99 b and particularly to a sub-cluster that contains strains from Nigeria alongside one strain from the Philippines (Figure 5). These two EV-C99 strains seem to be related to other strains recovered in 2015 from two children with AFP in Nigeria and another strain recovered in 2014 from a healthy child in the same country (Figure 5). The third NW-Nigeria EV-C99 strain recovered in this study belong to cluster EV-C99 c and is closely related to a strain recovered from children in Cameroon and Nigeria (Figure 5).

## DISCUSSION

In this study, we investigated whether the preponderance of EVs in AFP samples declared to be EV free by the RD-L20B cell culture based algorithm vary by region in Nigeria. Our results suggest that this might be the case. For example, while EVs were detected and unambiguously identified in 37.04% of the samples from NW Nigeria, they were only detected in 14.81% of the samples from SS Nigeria (Tables 2 and 3). Hence, for every 10 EVs found in such samples from SS Nigeria, 25 were found in an equivalent number of samples from NW Nigeria. The findings of this study are therefore in agreement with the history of varying regional preponderance and dynamics of enteroviruses (Faleye and Adeniji, 2015a) and polioviruses (CDC, 2006&2010; Mach et al., 2014; Duintjer-Tebbens et al., 2015) in Nigeria.

In this study, we also found that 78.57% (11/14) of the EVs detected were EV-Cs. This further confirms previous report (Oyero et al., 2014; Adeniji et al., 2017, Faleye et al., 2017) on EV-B bias of the RD component of the RD-L20B cell culture based algorithm (WHO, 2003, 2004). This allows selective accumulation as ‘negatives’, AFP samples that lacked EV-Bs but possibly had EV-Cs. Furthermore, our results suggest that the preponderance of EV-Cs in AFP samples declared to be EV free by the RD-L20B cell culture based algorithm vary by region in Nigeria. For instance, while EV-Cs were found in 29.6% (8/27) of the samples from NW Nigeria, they were only detected in 11.1% (3/27) of the samples from SS Nigeria (Tables 2 and 3). Consequently, for every 10 EV-Cs found in AFP samples declared to be EV free by the RD-L20B cell culture based algorithm from SS Nigeria, we have 27 in an equivalent number and type of samples from NW Nigeria.

It is known that the population gut (mucosal) immunity to PVs significantly determines the proportion of such population that can participate in PV transmission and consequently, the duration of transmission (Nathanson and Kew, 2010; Wassilak et al., 2011). However, it appears the higher preponderance of EV-Cs in NW Nigeria compared to SS Nigeria might have also contributed to the emergence in NW of over 50% (12/23) of the lineages of cVDPV2 (most of which were recombinants with NPEV-C non-structural region) that circulated in Nigeria between 2005 and 2011(Burns et al., 2013). The results of this study therefore, further highlight the importance of NPEV-Cs and the need to carefully and exhaustively catalogue their diversity and geographical distribution in Nigeria (considering their role in VDPV emergence [Burns et al., 2013]).

In this study, we detected CV-A1, CV-A19 and EV-C116 (Figure 2); enterovirus types whose prototype strains have not been grown in cell culture (Schmidt et al., 1975; Lipson et al., 1988; Lukashev et al., 2012). Particularly, they are known not to grow in RD cell culture (Lukashev et al., 2012; Tokarz et al., 2013). Therefore, the selective accumulation of these EV-C types in AFP samples that showed no growth in RD and L20B cell lines is in conformation with previous studies (Schmidt et al., 1975; Lipson et al., 1988; Lukashev et al., 2012; Tokarz et al., 2013). More recently, Sun et al., (2012) described two CV-A1 strains that developed cytopathology in RD cell culture. In 2015, we also (Adeniji et al., unpublished) recovered a CV-A1 strain that developed cytopathology in RD cell culture (Figure 2). Particularly interesting is the fact that this CV-A1 strain is very similar to the strain recovered in this study, despite the fact that it was recovered from NW-Nigeria about one month before the strain recovered in this study which was from SS-Nigeria (Figure 2). Why the NW-Nigerian strain replicated in cell culture but not the SS-Nigerian strain (despite their similarity in the VP1 region) is not clear. One possibility is the likely possession of different non-structural regions which might be responsible for the varying phenotypes as have been described for the polioviruses (Combellas et al., 2011). One other striking observation about the NW-Nigerian CV-A1 strain (Adeniji et al., unpublished) is the fact that it was growing simultaneously in RD cell culture alongside a CV-B4. Though only the CV-A1 virus was detected in the stool suspension, both CV-A1 and CV-B4 were detected in RD cell culture isolate. Thereby suggesting the CV-B4 strain might have been at very low titre in the faecal suspension. Furthermore, considering that in our laboratory, the two times we have detected CV-A24 in RD cell culture till date, it was replicating alongside Echovirus 7 (E7) in one instance and CV-B6 in the other (unpublished data), it therefore seems that co-replication with a CPE producing specie B enterovirus appears to either enable some EV-C members replicate in RD cell culture or facilitate their detection. However, more extensive investigations might be required to determine whether these observations are just chance occurrences or have a biological basis.

To the best of our knowledge, this represents the first description of Nigerian strains of CV-A19 and EV-C116. However, while the CV-A19 strain was easily identified unambiguously, identifying the EV-C116 was challenging. The EV strain identified as EV-C116 using the EGT phylogenetic tree (supplementary figure 1) did not cluster with the other EV-C116 strains recovered from GenBank (Figure 2). Rather it clustered with the reference CV-A22 strain Chulman (AF499643) and two other CV-A22 strains (JN542510 and KF412961) (Figure 2). When we further subjected the two non-Chulman CV-A22 strains (JN542510 and KF412961) that clustered with the EV-C116 strain identified in this study, to the EGT, they both typed as EV-C116. This suggests that these CV-A22 strains (JN542510 and KF412961) were really not CV-A22 but erroneously typed as such based on the sequence data available in GenBank as at the time they were initially sequenced. Puzzling, however, is the fact that the Chulman strain CV-A22 clustered with this group (CV-A22/EV-C116? [Figure 2]) but not with the remaining CV-A22 sequences (Figure 2) and the Oberste et al., (1999) enterovirus classification criteria (Table 4) validates the phylogenetic analysis results. Considering the strong bootstrap support both clusters have (Figure 2), the question that begs answering is, which of the two clusters is CV-A22 and what is the identity of the other?

The CV-A10 described in this study is the first strain of this EV type to be described in Nigeria, to the best of our knowledge. CV-A10, alongside EV-A71 and other EV-A types have been implicated in hand, foot and mouth disease (HFMD) (Tian et al., 2014; Tao et al., 2017) which sometimes might have neurological complications.Though CV-A10 is being described for the first time in Nigeria, EV-A71 has been detected repeatedly (Oyero et al., 2014; Faleye et al., 2016). Usually, EV-A71 strains (of genotype E) which are not associated with HFMD are detected in Nigeria (Oyero et al., 2014; Faleye et al., 2016). However, as Fernandez-Garcia et al., (2016) found in some West-African countries in 2013 and 2014, we also found in 2014 (unpublished data), a genogroup C of EV-A71 (which is associated with HFMD) in a child with AFP but co-infected with Echovirus 13 (E13). These sightings of CV-A10 and EV-A71 genogroup C in children with AFP in Nigeria might be a clarion call for a more intensive search for HFMD in the population or better still, a call to ascertain the reason the disease condition is hard to find in the population in spite of the presence of these viruses. It is however crucial to mention that as shown for the other EV types phylogenetically analyzed in this study (Figures 2-5), it is likely that several lineages of CV-A10 and EV-A71 genogroup C are present and circulating in Nigeria. Hence, the sporadic detection of these viruses in the country should not be interpreted as evidence of their absence. Rather, it should be appropriately seen as detection of the tip of the iceberg and consequently, incentive for surveillance of these viruses in the population.

Finally, in this study we were able to detect EVs in 29.63% (16/54) of the RD-L20B cell culture negative AFP samples from children <15 years old screened. We were however only able to identify the EVs present in 25.93% (14/54) of the samples. Consequently, as found in this study, the preponderance of EVs in such samples is lower than the 46.7% (14/30) previously described (Adeniji et al., 2017). Considering the samples analyzed in this study and those described in Adeniji et al., 2017 were all recovered from AFP cases in August 2015 (but independently and without replacement), it means that in all, 33.33% (28/84) of the samples had EVs in them. This value (33.33%) might therefore be a more accurate description of the prevalence of EVs in RD-L20B cell culture negative AFP samples from children <15 years old in Nigeria. It is however likely that the more samples of this type analyzed, the better we might get at reproducibly estimating the true prevalence of EVs in such sample types.

Of note, we found a Sabin strain PV2 in Adeniji et al., 2017 but found none in this study. This implies that only 1.2% (1/84) of the samples analyzed so far had poliovirus present. It however, clearly shows the downside of analyzing this sample set by random sampling especially considering the significance of missing poliovirus strains present in such to the eradication programme. Against this backdrop, it is essential that all (and not a random sample) of AFP samples declared negative for EVs by RD-L20B cell culture be further screened using cell culture independent techniques. Again, the need to ensure that no circulating PV strain is missed irrespective of whether or not it is from a difficult-to-access geographical location is paramount to the eradication programme. Hence, cannot be overemphasized.

## COMPLIANCE WITH ETHICAL STANDARDS

### Ethical Approval

There was no contact with human participants by any of the authors, and the article does not contain any information that can be used to associate the strains analyzed in this study to any individual.

### Funding

This study was funded by contributions from the authors

## ACKNOWLEDGEMENTS

We thank the WHO National Polio Laboratory in Ibadan, Nigeria for providing the anonymous samples analyzed in this study.

## AUTHOR CONTRIBUTIONS

1. Study Design (DE, JAA, MOA, TOCF)
2. Sample Collection, laboratory and Data analysis (All authors)
3. Wrote, revised, read and approved the final draft of the Manuscript (All authors)

## REFERENCES

1. Adeniji J.A. and Faleye, T.O.C. (2014b). Impact of Cell Lines Included in Enterovirus Isolation Protocol on Perception of Nonpolio Enterovirus Species C Diversity. Journal of Virological Methods. 10.1016/j.jviromet.2014.07.016

2. Adeniji, J. A. and T.O.C. Faleye (2014a). Isolationandidentificationofenterovirusesfrom sewageandsewagecontaminatedwater inLagos,Nigeria.FoodandEnvironmental Virology, 2014a, 6:75–86

3. Adeniji, J.A., Oragwa, A.O., George, U.E. Ibok, U.I., Faleye, T.O.C. Adewumi, M.O. (2017) Preponderance of Enterovirus Species C in RD-L20B cell culture negative stool samples from children diagnosed with Acute Flaccid Paralysis in Nigeria. Archives of Virology. DOI 10.1007/s00705-017-3466-2.

4. Burns, C.C., Diop, O.M., Sutter, R.W., Kew, O.M. Vaccine Derived Polioviruses. Journal of Infectious Disease. 2014; 210 (Suppl 1): S283–293.

5. Burns, C.C., J. Shaw, J. Jorba, et al., (2013). Multiple independent emergences of type 2 vaccine-derived polioviruses during a large outbreak in northern Nigeria, Journal of Virology, vol. 87, no. 9, pp. 4907–22. 2013.

6. Centers for Disease Control and Prevention. Outbreaks following wild poliovirus importations—Europe, Africa, and Asia, January 2009-September 2010. MMWR Morb Mortal Wkly Rep. 2010;59(43):1393–9. Epub 2010/11/05. mm5943a1 [pii]. pmid:21048560

7. Centers for Disease Control and Prevention. Resurgence of wild poliovirus type 1 transmission and consequences of importation—21 previously polio-free countries, 2002–2005. MMWR Morb Mortal Wkly Rep. 2006;55(6):145–50. pmid:16484977

8. Combelas N., Holmblat B., Joffret M. L., Colbère-Garapin F., Delpeyroux F. (2011). Recombination between poliovirus and coxsackie A viruses of species C: a model of viral genetic plasticity and emergence. Viruses 3, 1460–1484

9. DuintjerTebbens RJ, Pallansch MA, Wassilak SGF, Cochi SL, Thompson KM (2015). Combinations of Quality and Frequency of Immunization Activities to Stop and Prevent Poliovirus Transmission in the High-Risk Area of Northwest Nigeria. PLoS ONE 10(6): e0130123. doi:10.1371/journal.pone.0130123

10. Faleye T.O.C. and Adeniji J.A.(2015a) Enterovirus Species B Bias of RD Cell Line and Its Influence on Enterovirus Diversity Landscape. Food and Environmental Virology. 2015a; 7(4): 390–402.

11. Faleye, T.O.C., and Adeniji, J.A. (2015b) Nonpolio enterovirus-C (NPEV-C) strains circulating in South-Western Nigeria and their contribution to the emergence of recombinant cVDPV2 lineages. Brit. J. Virol. 2(5): 68–73.

12. Faleye, T.O.C., Adewumi, M.O., Coker, B.A., Nudamajo, F.Y. & Adeniji J.A. (2016). Direct Detection and Identification of Enteroviruses from Faeces of Healthy Nigerian Children Using a Cell-Culture Independent RT-Seminested PCR Assay. Advances in Virology, Article ID: 1412838, 12 pages, ttp://dx.doi.org/10.1155/2016/1412838.

13. Faleye TOC, Adewumi MO, Japhet MO, David OM, Oluyege AO, Adeniji JA, Famurewa O. (2017). Non-polio enteroviruses in faeces of children diagnosed with acute flaccid paralysis in Nigeria. Virology Journal, 14:175, DOI 10.1186/s12985-017-0846-x

14. Fernandez-Garcia MD, Kebe O, Fall AD, Dia H, Diop OM, Delpeyroux F, Ndiaye K. (2016). Enterovirus A71 Genogroups C and E in Children with Acute Flaccid Paralysis, West Africa.Emerg Infect Dis. 2016 Apr;22(4):753–5. doi: 10.3201/eid2204.151588.

15. Kimura, M. (1980). Asimplemethodforestimatingevolutionaryrateofbasesubstitutions through comparative studiesof nucleotide sequences. JournalofMolecular Evolution. 16(2): 111–120.

16. Kroneman, A., Vennema, H., Deforche, K., Van der Avoort, H., Penarandac, S., Oberste, M.S., Vinjéc, J., Koopmans, M., (2011). An automated genotyping tool for enteroviruses and noroviruses. Journal of Clinical Virology. 51, 121–125.

17. Lipson S. M., Walderman R., Costello P., Szabo K. (1988). Sensitivity of rhabdomyosarcoma and guinea pig embryo cell cultures to field isolates of difficult-to-cultivate group A coxsackieviruses. J ClinMicrobiol 26, 1298–1303

18. Lukashev A. N., Drexler J. F., Kotova V. O., Amjaga E. N., Reznik V. I., Gmyl A. P., Grard G., TatyTaty R., Trotsenko O. E. & other authors (2012). Novel serotypes 105 and 116 are members of distinct subgroups of human enterovirus C. J Gen Virol 93, 2357–2362.

19. Mach O, Tangermann RH, Wassilak SG, Singh S, Sutter RW. Outbreaks of paralytic poliomyelitis during 1996–2012: the changing epidemiology of a disease in the final stages of eradication. J Infect Dis. 2014;210(Suppl 1):S275–S82. pmid:25316846

20. Nathanson, N. and Kew, O. M. (2010). From Emergence to Eradication: The Epidemiology of Poliomyelitis Deconstructed. American Journal of Epidemiology. 172(11), 1213 – 1229.

21. Nix, W. A., Oberste, M. S., Pallansch, M. A. (2006) Sensitive, Seminested PCR Amplification of VP1 Sequences for Direct Identification of All Enterovirus Serotypes from Original Clinical Specimens. Journal of Clinical Microbiology, 44(8): 2698–2704

22. Oberste, M. S., Maher, K., Kilpatrick, D. R., Pallansch, M. A. (1999). Molecular evolution of the human enteroviruses: correlation of serotype with VP1 sequence and application to picornavirus classification. Journal of Virology. 73(3), 1941–1948.

23. Oyero, O. G., F. D. Adu, J. A. Ayukekbong. (2014). Molecular characterization of diverse species enterovirus-B types from children with acute flaccid paralysis and asymptomatic children in Nigeria. Virus Research. 189:189–193

24. Pipkin, P.A., Wood, D.J., Racaniello, V.R., Minor, P.D. Characterization of L cells expressing the human poliovirus receptor for the specific detection of polioviruses in vitro. Journal of Virological Methods. 1993; 41: 333–340.

25. Sadeuh-Mba, S. A., Bessaud, M., Massenet, D., Joffret, M. L., Endegue, M. C., Njouom, R., Reynes, J. M., Rousset, D., Delpeyroux, F., (2013). High frequency and diversity of species C enteroviruses in Cameroon and neighboring countries. Journal of Clinical Microbiology. 51, 759–770.

26. Schmidt N. J., Ho H. H., Lennette E. H. (1975). Propagation and isolation of group A coxsackieviruses in RD cells. J ClinMicrobiol 2, 183–185.

27. Sun Q., Zhang Y., Zhu S., Cui H., Tian H., Yan D., Huang G., Zhu Z., Wang D. & other authors (2012). Complete genome sequence of two coxsackievirus A1 strains that were cytotoxic to human rhabdomyosarcoma cells. J Virol 86, 10228–10229.

28. Tamura, K., Peterson, D., Peterson, N., Stecher, G., Nei, M., Kumar, S., (2011).. MEGA5: molecular evolutionary genetics analysis using maximum likelihood, evolutionary distance, and maximum parsimony methods. Molecular Biology and Evolution. 28, 2731–2739.

29. Tao, J. et al. (2017). Epidemiology of 45,616 suspect cases of Hand, Foot and Mouth Disease in Chongqing, China, 2011-2015. Sci. Rep. 7, 45630; doi: 10.1038/srep45630.

30. Tian H, Zhang Y, Sun Q, Zhu S, Li X, et al. (2014) Prevalence of Multiple Enteroviruses Associated with Hand, Foot, and Mouth Disease in Shijiazhuang City, Hebei Province, China: Outbreaks of Coxsackieviruses A10 and B3. PLoS ONE 9(1): e84233. doi:10.1371/journal.pone.0084233

31. Tokarz R., Hirschberg D. L., Sameroff S., Haq S., Luna G., Bennett A. J., Silva M., Leguia M., Kasper M. & other authors (2013). Genomic analysis of two novel human enterovirus C genotypes found in respiratory samples from Peru. J Gen Virol 94, 120–127.

32. Wassilak S1, Pate MA, Wannemuehler K, Jenks J, Burns C, Chenoweth P, Abanida EA, Adu F, Baba M, Gasasira A, Iber J, Mkanda P, Williams AJ, Shaw J, Pallansch M, Kew O. (2011) Outbreak of type 2 vaccine-derived poliovirus in Nigeria: emergence and widespread circulation in an underimmunized population. J Infect Dis. 203(7):898–909. doi: 10.1093/infdis/jiq140.

33. World Health Organisation (1988). Global eradication of poliomyelitis by the year 2000 (World Health Assembly resolutionWHA41–28). http://www.polioeradication.org/content/publications/19880513resolution.pdf

34. World Health Organisation (2003). Guidelines for environmental surveillance of poliovirus circulation. Geneva.

35. World Health Organisation. (2004). Polio laboratoryManual. 4th Edition, Geneva.

36. World Health Organisation. (2015). Enterovirus surveillance guidelines: Guidelines for enterovirus surveillance in support of the Polio Eradication Initiative, Geneva.

37. www.picornaviridae.com

38. www.polioeradication.org

